# Spatial and temporal characterization of the rich fraction of plastid DNA present in the nuclear genome of *Moringa oleifera* reveals unanticipated complexity in NUPTś formation

**DOI:** 10.1101/2023.05.31.542793

**Authors:** Juan Pablo Marczuk-Rojas, Angélica María Álamo-Sierra, Antonio Salmerón, Alfredo Alcayde, Viktor Isanbaev, Lorenzo Carretero-Paulet

## Abstract

Beyond the massive amounts of DNA and genes transferred from the protoorganelle genome to the nucleus during the endosymbiotic event that gave rise to the plastids, stretches of plastid DNA of varying size are still being copied and relocated to the nuclear genome in a process that is ongoing and does not result in the concomitant shrinking of the plastid genome. As a result, plant nuclear genomes are featured by a small, but variable, fraction of their genomes of plastid origin, the so-called nuclear plastid DNA sequences (NUPTs). However, research on the topic was mostly focused on a limited number of species and of plastid DNA. Here, we leveraged a chromosome-scale version of the *Moringa oleifera* genome, featured by the largest fraction of plastid DNA in any plant nuclear genome, to examine the chromosomal distribution and arrangement of NUPTs, to explicitly model and test the correlation between their age and size distribution, as well as to characterize their sites of origin at the chloroplast genome and their sites of insertion at the nuclear one. We found a bimodal distribution of NUPT relative ages, which implies NUPTs in moringa were formed through two separate events. Furthermore, NUPTs from every event showed markedly distinctive features, suggesting they originated through distinct mechanisms. Our results reveal an unanticipated complexity of the mechanisms at the origin of NUPTs and of the evolutionary forces behind their fixation and highlight moringa species as an exceptional model to study the mechanisms of origin and evolutionary fate of NUPTs.

**Significance statement:** Sequencing of plant genomes has revealed the presence of a variable, but significant, content of DNA arising from their organelles, *i.e.*, plastids and mitochondria. However, the mechanisms underlying their origin and fixation are not yet fully elucidated. Here, by thoroughly examining the spatial and temporal distribution of the large fraction of plastid DNA present in the nuclear genome of the orphan crop *Moringa oleifera*, we identified their episodic origin trough seemingly distinct mechanisms, highlighting the complexity of the molecular and evolutionary dynamics underlying their origin and fixation and revealing moringa as an outstanding model to assess the impact of plastid DNA in the evolution of the architecture and function of plant nuclear genomes.

## Introduction

Nearly all plant nuclear genomes contain a small, but significant, fraction of their nuclear genomes composed of DNA sequences derived from their chloroplasts (Zhang et al. 2020); these nuclear integrants of plastid DNA are commonly known as nuclear plastid DNA sequences (NUPTs) (Timmis et al. 2004). The process of NUPTś formation has been commonly associated to the process by which most genes present in the bacterial ancestor of plastids were transferred to the nuclear genome and their products eventually retargeted to their ancestral compartment during the endosymbiotic event that gave rise to the chloroplast organelle. However, whereas the latter entails the loss of vast amounts of DNA with the subsequent reduction of its size and the transfer of most of the genes originally present in the protoorganelle organism to the nuclear genome (Martin et al. 1998, 2002), the former involves the copy of stretches of DNA from the chloroplast genome. Even though most NUPTs are less than 1 kb in length, NUPTs of recent origin spanning the whole chloroplast chromosome have been detected in *Arabidopsis thaliana* (Arabidopsis)*, Oryza sativa* (rice) and *Populus trichocarpa* (Huang et al. 2005, 2017), and did not result in the shrinking of the plastid genome.

Although the process of NUPTs’ formation is still poorly understood, it is expected to involve the following sequence of events. First, the duplication of a stretch of DNA present in the chloroplast genome. Second, the lysis of chloroplast organelle membranes to allow the leakage of duplicated plastid DNA. Third, the import to the nucleus of the leaked plastid DNA. Fourth, the integration of plastid DNA into the nuclear genome. At present, no mechanism has been formally proposed to explain the recurrent duplication of stretches of plastid DNA of varying sizes that are at the origin of NUPTs. The biological mechanisms involved in the leakage of plastid DNA to the cytoplasm and its subsequent import by the nucleus are not yet completely elucidated either, although gametogenesis and cell stress (especially pollen development and mild heat stress, respectively) have been reported to induce the disruption of chloroplast organelle membranes (Richly & Leister 2004; Timmis et al. 2004; Sheppard et al. 2008; Kleine et al. 2009; Wang, et al. 2012). It has been also suggested that certain kinds of stresses, such as ionizing radiation and pathogen infections, may, not only trigger the leakage of plastid DNA to the nucleocytosolic compartment, but also favor its integration into the nuclear genome (Bock & Timmis 2008). The molecular mechanisms of NUPTś integration into the nuclear genome are not fully described either, but they are probably diverse and generally involve double-stranded breaks (DSBs) and DNA damage and thus are potentially mutagenic. For example, it has been hypothesized that NUPTś integration is mediated by non-homologous end joining (NHEJ) during DSB repair events (Leister & Kleine 2011; Wang & Timmis 2013; Wang et al. 2018). Another suggested mechanism is homologous recombination via gene conversion or single strand annealing (SSA), another DSB repair pathway (Portugez et al. 2018). Furthermore, integration events can involve one or multiple plastid DNA sequences which, in turn, originate continuous NUPTs or NUPT mosaics, respectively (Huang et al. 2004; Noutsos et al. 2005; Portugez et al. 2018; Wang et al. 2018).

Because of i) the potential mutagenic events of NUPTs’ formation, and ii) the uncontrolled proliferative insertion of organelle DNA might lead to the unnecessary “obesity” of the nuclear genome (Kleine et al. 2009; Zhang et al. 2020), most NUPTs are expected to be rapidly fragmented and shuffled away through transpositions and genome arrangements and, eventually, purged from the nuclear genome (Matsuo et al. 2005; Michalovova et al. 2013; Yoshida et al. 2014). As a consequence, the distribution of NUPTs by age should follow an exponential distribution, indicating a continuous rate of NUPTs’ formation and decay throughout time (Matsuo et al. 2005). Although such a pattern has been suggested for rice, *Medicago truncatula*, *P. trichocarpa* and *Zea mays* (Matsuo et al. 2005; Chen et al. 2015; Yoshida et al. 2014), different patterns have been observed in other species such as Arabidopsis, *Carica papaya*, *Fragaria vesca*, *Moringa oleifera* and *Vitis vinifera* (Yoshida et al. 2014; Chen et al. 2015; Ojeda-López et al. 2020). A second consequence is the expected positive correlation between NUPTs’ size and age, an observation that has been suggested for several species, despite not being explicitly tested statistically (Richly & Leister 2004; Michalovova et al. 2013; Yoshida et al. 2014; Li et al. 2019).

Indeed, the fraction of nuclear genomes occupied by NUPTs varies enormously among species and even within different populations of the same species (Huang et al. 2005; Roark et al. 2010; Ma et al. 2020). Most species showed around 0.1% of plastid DNA in their nuclear genome, with very few showing more than 1% (Zhang et al. 2020). These large variations in the fraction of nuclear genomes occupied by NUPTs raise the question of what evolutionary forces may lie behind the fixation of variable fractions of plastid DNA in plant nuclear genomes. However, previous studies on the mechanisms of origin and evolutionary fate of NUPTs were mostly focused on a limited number of species and involved a reduced number of NUPTs. A more detailed picture will certainly benefit from a larger number of NUPTs and a higher fraction of the nuclear genome occupied by plastid DNA.

So far, the largest fraction of DNA of plastid origin found in any plant nuclear genome (4.71%) has been detected in the orphan crop *M. oleifera* (moringa) (Ojeda-López et al. 2020). In the present study, we leveraged a recent chromosome-scale version of the moringa genome (Chang et al. 2022) to examine the spatial distribution and arrangement in clusters of NUPTs, to explicitly model and test the correlation between their age and size distribution, as well as to characterize their sites of origin at the chloroplast genome and their sites of insertion at the nuclear one. Our results reveal an unanticipated complexity of the mechanisms at the origin of NUPTs as well as the evolutionary forces behind their fixation.

## Results

### Widespread distribution of NUPTs in the moringa nuclear genome

In order to detect NUPTs present in the moringa nuclear genome, a chromosome-scale assembly of the moringa genome, AOCCv2 (Chang et al. 2022), was scanned using BLASTN and the moringa chloroplast genome sequence (Lin et al. 2019) as query. This resulted in 13,901 total alignments, which were defined as NUPTs in our analysis (**Supplementary Table S1**). 13 out of the 14 chromosomes hosted more than 100 NUPTs (ranging from 555 to 2,176) and only eight chromosomes and one scaffold contained NUPTs summing up above 160,600 bp (*i.e*., the size of the moringa chloroplast genome) (**Supplementary Table S1**).

The total aligned region between the chloroplast genome and the nuclear genome summed up a total of 11,286,242 bp, which represents a 4.77% of the size of the nuclear genome assembly, pretty similar to previous estimations of 4.71% obtained with a previous version of the moringa genome, AOCCv1 (Ojeda-López et al. 2020). After correcting for redundancy in BLASTN hits resulting from Inverted Repeat (IR) regions of the moringa chloroplast genome (1,276), the fraction of the moringa nuclear genome corresponding to NUPTs was of 3.92%. As a comparison, the same scanning procedure was applied to the nuclear genomes of Arabidopsis and rice (Kawahara et al. 2013; Cheng et al. 2017) (Table 1) using their respective chloroplast genomes (Hiratsuka et al. 1989; Sato et al. 1999) (Table 1) as queries; the fraction of plastid DNA was 0.083% and 0.34% (0.073% and 0.28% after correcting for redundant hits involving IR regions), respectively, close to previous estimates (Richly & Leister 2004; Zhang et al. 2020).

**Table 1:** Summary of genome versions used in this study.

| **Species** | **Nuclear genome**<br>**version** | **Chloroplast genome version**<br>**(NCBI RefSeq number)** |
| --- | --- | --- |
| *Moringa oleifera* | AOCC v2 | NC_041432.1 |
| *Arabidopsis thaliana* | TAIR Araport11 | NC_000932.1 |
| *Oryza sativa ssp. japonica* | IRGSP-1.0 | NC_001320.1 |

### Most NUPTs in moringa originated through two distinct events

In order to gain insights on the timing of plastid DNA acquisition by the moringa nuclear genome, we examined the relative age distribution of NUPTs using the percent identity of the corresponding BLASTN hits as a proxy of evolutionary time. Assuming the mutation rate is proportional to evolutionary time*, i. e*., the molecular clock hypothesis holds, the lower the percent identity, the older the NUPTs. Percent identity of BLASTN hits ranges between 71.43 and 100% and shows an apparent bimodal distribution (Figure 1A). Indeed, when Gaussian mixture models were fitted to the corresponding density curves, two clear peaks, centered around 74.71 and 92.52%, respectively, were detected (Figure 1A). According to the posterior probabilities of assigning a NUPT to either one or another peak, using a threshold of 95%, 8,388 NUPTs (60.34% of the total) summing up a total of 1,534,130 bp (13.59% of the total) belonged to the older peak (from now on Episode I, or NUPTs-I), while 4,530 NUPTs (32.59% of the total) summing up a total of 9,481,497 bp (84.01% of the total) belonged to the younger peak (from now on Episode II or NUPTs-II). The rest of NUPTs (983) were not confidently assigned to either one or the other peak. Taking as a whole, these results support two main episodic formation events at the origin of most NUPTs.

**Fig. 1.**
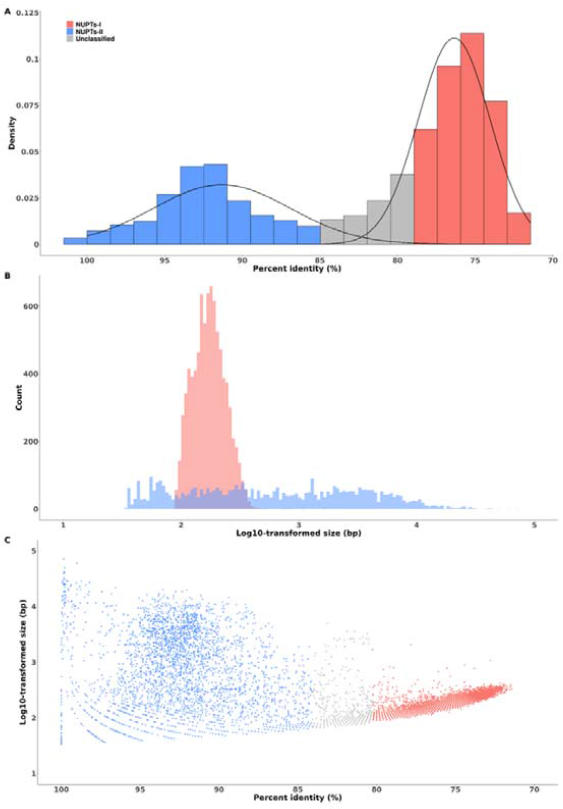
Modeling the distribution of percent identity and size of moringa NUPTs. A, histogram of the distribution of NUPTs percent identity values. The two density plots resulting from fitting Gaussian mixture models, putatively corresponding to distinct events of NUPTs formation (I and II), are shown. B, Histogram of the distribution of NUPTs size values. C, scatterplot of percent identity versus sizes of moringa NUPTs. For an easier visualization, NUPT size values have been log 10-transformed.

Next, we examined the size distribution of NUPTs, partitioned by each of the retrieved episodes. While NUPTs-I ranged in size from 89 to 2,048 bp, NUPTs-II ranged from 33 to 71,935 bp (Figure 1B). Both followed a non-normal right-skewed unimodal distribution (Figure 1B), with a mean and a median size of 182.9 and 172 or 2,093.05 and 550.5 bp for NUPTs-I and NUPTs-II, respectively.

From studies in rice and other plant species, it had been suggested an apparent positive correlation between size and sequence identity of NUPTs, *i.e*., larger NUPTs tend to be more conserved at the sequence level. This observation can be interpreted as young, larger conserved NUPTs declining and fragmenting over time, and eventually being purged from the genome (Richly & Leister 2004; Noutsos et al. 2005; Matsuo et al. 2005; Michalovova et al. 2013; Yoshida et al. 2014; Li et al. 2019). To test whether this observation also applied to moringa NUPTs, we studied the correlation between size and sequence identity by means of two different tests accounting for not-normally distributed data, again partitioned by every episode detected (Figure 1C and Table 2). Interestingly, while for younger NUPTs from episode II size positively correlated with sequence identity in both tests (Table 2), the opposite trend was observed for NUPTs-I (Table 2), suggesting different mechanisms might have been at the origin of NUPTs from every episode and / or, once integrated, they might also have followed different evolutionary trajectories.

**Table 2:** Correlation analysis between NUPTs’ sequence identity and size by NUPTs’ formation event.

| **Method** | **Correlation**<br>**coefficient** | ***P*-value** | **Episode** |
| --- | --- | --- | --- |
| Kendall's rank correlation tau | -0,71 | $< 2.2 \times 10^{-160}$ | I |
| Spearman's rank correlation rho | -0,86 | $< 2.2 \times 10^{-160}$ | I |
| Kendall's rank correlation tau | 0,07 | $4.59 \times 10^{-6}$ | II |
| Spearman's rank correlation rho | 0,10 | $1.31 \times 10^{-4}$ | II |

### Characterization of the differential distribution of NUPTś insertion sites in the moringa nuclear genome

The distribution of NUPTs across the 14 chromosomes conforming the moringa nuclear genome was represented in a Circos plot as independent density plots for every episode (Figure 2). In contrast to NUPTs-I, many NUPTs-II appeared to be highly concentrated in some specific regions of chromosomes one, five, six and 10 which showed prominent peaks in the density plots, likely corresponding to hotspots where NUPTs integration and / or subsequent fixation is favored (Figure 2).

**Fig. 2.**
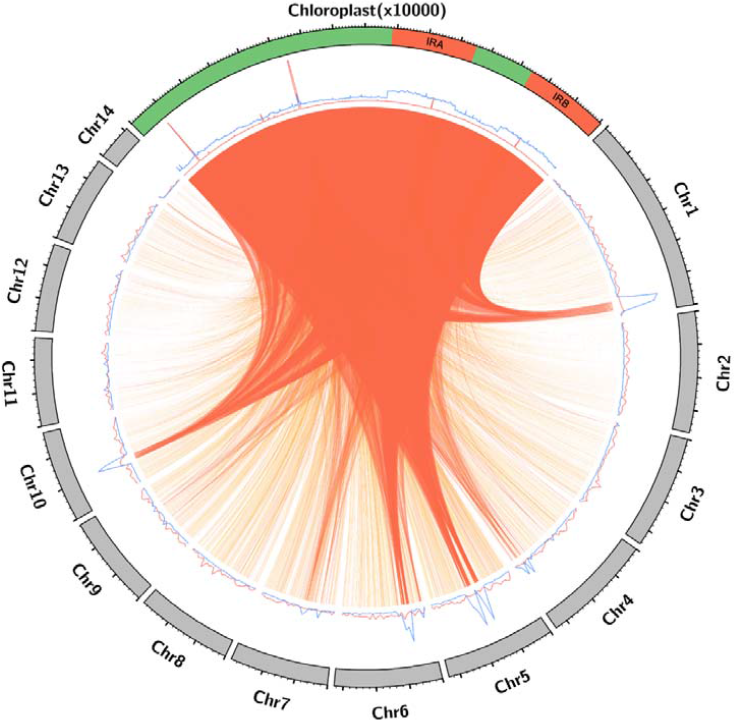
Circos plot representation of NUPTs in the moringa nuclear genome. Nuclear chloroplast and chromosomes are represented as grey and green filled blocks, respectively, forming a circumference. Red bands in the chloroplast genome represent Inverted Repeat (IR) regions. Results are shown for the 14 nuclear chromosomes, hosting 13,352 NUPTs representing a total of 10,406,471 bp, *i.e.*, 96.05% and 92.20% of the total, respectively. The block corresponding to the chloroplast genome is located at 12 o’clock, and the 14 nuclear chromosomes are arranged clockwise. Nuclear chromosomes are drawn to scale, with lengths proportional to size and expressed in Mb, while the chloroplast genome has been upscaled to occupy a quarter of the image circumference; its size unit was set to 10,000 bp. Line plots representing the respective density distributions of NUPTs-I (red) and NUPTs-II (blue) are displayed. Windows of 500,000 and 100 bp were selected for the nuclear and chloroplast chromosomes, respectively. Local BLASTN sequence alignments between the chloroplast and the nuclear genome corresponding to putative NUPTs are represented as ribbons. Ribbons are colored according to the percentage of sequence identity of the local alignments (NUPTs) grouped by quartiles (with yellow, light orange, orange, and red corresponding to the first, second, third and fourth quartiles, respectively).

A recent survey in African and Asian rice reported a compositional bias at the flanking regions of NUPTs’ insertion sites (Ma et al. 2020). A similar compositional bias was actually observed in moringa, *i.e*., the 100 bp regions flanking NUPT-I and NUPT-II displayed a lower GC content on average (17.63 and 31.84%, respectively) than the rest of the genome (35.9%), after excluding NUPT sequences, with differences being significant according to Mann-Whitney U-tests (*P* < 2.2 x 10^-260^; *P* = 3.83 x 10^-174^, respectively).

Previous analysis on NUPTs from Arabidopsis and rice identified their tendency to group in clusters, defined as a group of two or more non-overlapping NUPTs where the distance between two consecutive integrants was less than 5 kb (Richly & Leister 2004). We tried to determine whether NUPTs in moringa were also forming clusters. 3,135 NUPTs (22.55% of the total) summing up a total of 3,274,319 bp (29.01% of the total) were found grouping into 1,034 clusters, which were detected in the 14 chromosomes and six scaffolds, and whose sizes ranged from 122 to 110,808 bp (**Supplementary Tables S2**).

Then we examined separately clusters grouping NUPTs from every episode. 1,114 NUPTs-I (*i.e.*, 13.28%) summing up a total of 207,462 bp (*i.e.*, 13.52%) were found forming 504 clusters which hosted up to six integrants (Figure 3) (**Supplementary Table S2**), whereas 1,044 NUPTs-II (*i.e.*, 23.05%) summing up a total of 2,645,573 bp (*i.e.*, 27,90%) were found inside 259 clusters which hosted up to 21 integrants (Figure 3) (**Supplementary Table S2**). The rest of the clusters (271) hosted 977 NUPTs from either one or both episodes and / or unclassified NUPTs (**Supplementary Table S2**).

**Fig. 3.**
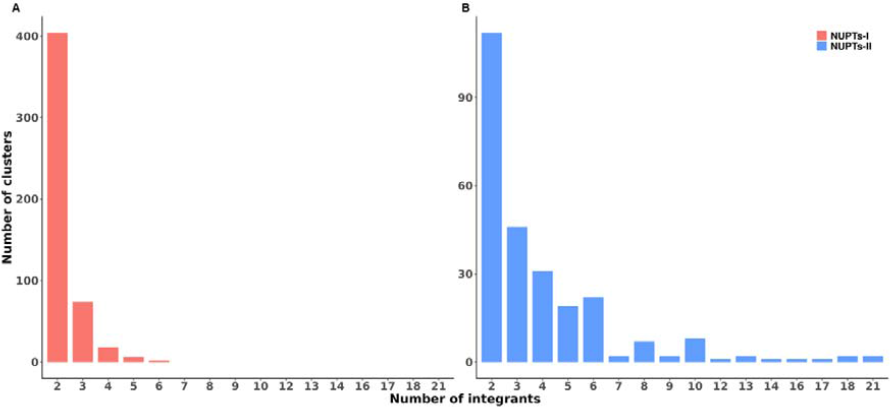
Distribution of the number of integrants of clusters of NUPTs.

The grouping into clusters of NUPTs at specific positions might be reflecting either large NUPTs fragmenting over time after their integration into the nuclear genome or chromosomal hotspots. If the former were the case, the sequence identity of NUPTs should correlate with their tendency to group into clusters. To test this hypothesis, we examined the correlation between the average sequence identity of the NUPTs in every cluster and the number of integrants. The tests were performed separately on clusters formed exclusively by NUPTs-I and NUPTs-II. While average sequence identity positively correlated with the number of NUPTs-II inside clusters in both correlation tests (Table 3), no significant correlation was found for NUPTs-I (Table 3). The different trends observed also supported different mechanisms at the origin of NUPTs from one or another episode, and eventually also their subsequent evolutionary fate.

**Table 3:** Correlation analysis between average sequence identity of NUPTs in every cluster and the number of integrants partitioned by episode of NUPTs’ formation.

| **Method** | **Correlation**<br>**coefficient** | ***P*-**<br>**value** | **Episode** |
| --- | --- | --- | --- |
| Kendall's rank correlation tau | -0.0461 | 0.197 | I |
| Spearman's rank correlation rho | -0.0575 | 0.198 | I |
| Kendall's rank correlation tau | 0.101 | 0.0267 | II |
| Spearman's rank correlation rho | 0.139 | 0.0252 | II |

### Biased distribution of NUPTs in the moringa chloroplast genome

Finally, we studied the distribution of NUPTs across the moringa chloroplast genome. For this purpose, we divided the corresponding DNA sequence into 100 bp regions. First, we performed the analysis using the whole set of NUPTs. Although all chloroplast genome regions were involved in forming NUPTs, two short regions were found to be involved in the majority of NUPTs. A 350 bp region spanning the interval 53,800-54,150 within the chloroplast genome was involved in more than half of the NUPTs (7,923, 57% of the total). A second 200 bp region spanning the interval 8,600-8,800 was involved in 734 NUPTs. These two regions have in common that they were mostly composed of A and T bases, consequently displaying an extremely low GC content (less than 0.05%) compared to the rest of the chloroplast genome (36.8%).

Finally, in order to examine whether these two regions contributed differentially to the formation of NUPTs-I and NUPTs-II, we subsequently performed this analysis separately for both sets. While the vast majority of the NUPTs-I (7,771, 92.64% of the total) originated from these two chloroplast genome regions (Figure 2), this was only true for only 332 NUPTs-II (7.33% of the total) (Figure 2). The rest of the NUPTs-II were almost uniformly distributed across the chloroplast genome, except for the IR regions, where, as expected, twice the number of NUPTS-II could be observed.

## Discussion

By leveraging a recently obtained high-quality long-read chromosome-scale assembly of the nuclear genome of moringa (*i.e.*, AOCCv2) (Chang et al. 2022), we gained a finer characterization of the rich fraction of plastid DNA originally detected in an older, less contiguous, version (*i.e.*, AOCCv1) (Chang et al. 2019), the highest reported for any plant species so far (Ojeda-López et al. 2020). While the total fraction of plastid DNA was similar using both versions of the genome, differences were observed regarding the events underlying such enrichment. Our previous report (Ojeda-López et al. 2020), using the distribution of synonymous substitutions rates as a proxy of evolutionary time, attributed such enrichment in plastid DNA to a recent single burst of plastid gene duplicates relocating to the moringa nuclear genome. Here, in turn, by fitting Gaussian mixture models to the distributions of sequence identity of NUPTs (taken instead as a proxy of evolutionary time), two distinct main episodic events of NUPTs’ formation could be detected, namely NUPTs-I and NUPTs-II. The reason for this discrepancy likely resides in errors in the annotation of the AOCCv1 moringa nuclear genome, featured by an overrepresentation of small genes annotated with chloroplast and photosynthetic functions. While 656 and 114 genes were annotated with the terms “chloroplast” or “photosynthesis”, respectively, in the AOCCv1 Moringa genome, only 378 and 51 genes were annotated with such terms in AOCCv2 (Chang et al. 2022). For example, while 45 fragmented nuclear genes were annotated as encoding for the plastid-encoded large subunit of ribulose-1,5-bisphosphate carboxylase/oxygenase (RBCL) in AOCCv1, only three were annotated as such in AOCCv2, although all of them could be mapped to specific genomic regions in AOCCv2. Altogether suggests the previous enrichment in chloroplast related functions observed among nuclear genes was likely due to fragmented DNA of plastid origin, *i.e*., NUPTs, encompassing coding regions, wrongly annotated as gene coding models.

Hitherto, relative ages of NUPTs’ formation in different plant species had been reported to be featured by either exponentially decreasing or uniformly constant distributions (Matsuo et al. 2005; Yoshida et al. 2014; Chen et al. 2015), which fit, respectively, into two different modes of NUPTś formation, *i.e.*, single events and hotspots (Richly & Leister 2004; Noutsos et al. 2005). The single event mode commonly results in long continuous NUPTs collinear with specific regions of the chloroplast genome, which are concentrated in specific regions of the nuclear genome, *e.g*., (peri)centromeric regions (Richly & Leister 2004; Noutsos et al. 2005; Matsuo et al. 2005; Michalovova et al. 2013), and are expected to decay into smaller fragments and relocate as a consequence of chromosomal rearrangements and reshuffling involving transposable element activity (Michalovova et al. 2013). In contrast, hotspots result in the concomitant integration of multiple short NUPTs from different origins arranged as a mosaic in specific loci of the nuclear genome (Huang et al. 2004; Noutsos et al. 2005).

To the best of our knowledge, no previous studies have reported the bimodal distribution of NUPT relative ages observed here for moringa, which implies NUPTs in moringa were formed through two events separated in time. Furthermore, NUPTs from every event showed markedly distinctive features, suggesting they originated through distinct mechanisms. For example, according to the relative distribution of sizes, younger NUPTs from episode II showed seemingly random origins throughout the chloroplast genome and were featured by a wide range of sizes, their preferential location in hotspots across the nuclear genome and a weak positive correlation between sequence identity and size. However, although some NUPTs-II may have originated as long fragments subsequently breaking into smaller pieces arranged collinearly as clusters throughout the nuclear genome, in accordance with the single event mode (Noutsos et al. 2005), a slight, but consistent positive correlation was observed between the number of NUPTs-II grouping in clusters and sequence identity. This correlation suggests at least some NUPTs-II may have also originated as smaller fragments landing in specific landmarks of the nuclear genome, *i.e.*, chromosomal hotspots, eventually further dispersing trough different kinds of genome rearrangements. Altogether supports the origin of NUPTs-II through both single events and hotspots modes

In turn, older NUPTs from episode I, featured by a narrower distribution of sizes and a significant negative correlation between sequence identity and size, do not seem to fit into any of the two modes of NUPTs’ formation previously described. Moreover, a vast majority of NUPTs-I originated from two short regions in the chloroplast genome, an observation only reported previously for *Asparagus officialis* (Li et al. 2019) and in contrast to previous studies in Arabidopsis, rice and other species, which showed a homogenous distribution of NUPTs throughout the chloroplast genome (Matsuo et al. 2005; Yoshida et al. 2014). We therefore propose here a third mode of NUPTs’ formation through small-scale recurrent events. We noted that both chloroplast regions at the origin of most NUPTs-I were featured by a strong enrichment in A and T bases forming repetitive sequences. Regions in the chloroplast genome containing A/T rich repetitive sequences have been reported as highly prone to inducing indels via replication slippage events (Massouh et al. 2016), which might provide a molecular mechanism for the duplication and leakage of specific regions of plastid DNA. Once individual NUPTs are formed, two scenarios are plausible i) multiple copies of NUPTs firstly forming in the chloroplast and later relocating to the nucleus, or ii) individual NUPTs recurrently duplicating once integrated into the nuclear genome.

Moreover, insertion sites for NUPTs-I and NUPTs-II throughout the nuclear genome are similarly enriched in A and T bases, which have been reported as fragile sites in which the formation of DNA secondary structures interrupting the replication bubble is favored, ultimately leading to DSB and, therefore, becoming potential suitable targets for promiscuous organellar DNA integration as suggested in (Ma et al. 2020). NUPTs’ insertion sites have also been reported to occur preferentially in open chromatin regions which, in turn, are also more exposed to proteins participating in DNA breakage and repair (Tsuji et al. 2012; Wang & Timmis 2013).

In respect of the possible evolutionary forces underlying the leakages and subsequent fixation of variable amounts of plastid DNA in plant nuclear genomes, these might be related to the different stressful conditions to which every species would have been subjected throughout their recent evolutionary history; different stresses have been shown to promote DNA migration from chloroplasts to the nucleus (Wang, et al. 2012; Cullis et al. 2009). For example, it has been noted that the 11 giant NUPTs found in Asian rice trended to distribute in natural populations from higher latitude regions featured by lower temperatures and light intensities (Ma et al. 2020). This observation led the authors to attribute NUPTs a potential role in enhancing environmental adaptation by increasing the number of chloroplast-derived genes which might, in turn, improve photosynthesis (Ma et al. 2020). Similarly, the massive amounts of plastid DNA found in the moringa nuclear genome might be related to the exposure to stressful conditions during its recent evolutionary history. Indeed, domestication of moringa from the sub-Himalayan lowlands in NW India, its putative location of origin where mean annual precipitations exceed 1100 mm, to tropical and sub-tropical areas around the world where its culture has spread (Pandey et al. 2011), would have likely involved the selection of varieties better adapted to drier and hotter environments (Brunetti et al. 2018, 2020). Indeed, moringa has been reported to successfully cope with multiple stresses, particularly water deficit and UVB radiation (Araújo et al. 2016). Comparative genomics of domesticated moringa together with that of the 12 wild Moringa species that make up the taxonomic family Moringaceae within the Brassicales order (Olson 2002), emerges as an excellent model for reconstructing the mechanisms of origin and evolutionary fixation of plastid DNA in the nuclear genome.

### Experimental procedures

#### Detection of plastid DNA in the nuclear genome

NUPTs in the nuclear genomes of moringa (Chang et al. 2022), Arabidopsis (Cheng et al. 2017) and rice (Kawahara et al. 2013) were detected using the BLASTN local alignment tool from the BLAST+ program package v2.12.0+ (Altschup et al. 1990). In each case, the chloroplast genome sequences of moringa (Lin et al. 2019), Arabidopsis (Sato et al. 1999) and rice (Hiratsuka et al. 1989) (Table 1) were used as queries and the corresponding nuclear genome sequences (Table 1) as databases. The parameters were as follows: `-evalue 1e-5-word_size 9 -penalty −2 -show_gis -dust no -num_threads 8`. Results in terms of sequence identity and density of NUPTs were represented as circular plots, constructed using Circos version 0.69−8 (Krzywinski et al. 2009). In order to correct for redundancy of NUPTs resulting from the inverted repeat (IR) region of the chloroplast genome, BLASTN hits were counted only once.

#### Gaussian mixture modeling of NUPTs’ percent identity distribution

In order to detect peaks in the distribution of percent identity values putatively corresponding to episodic events of NUPTs integration in the nuclear genome, Gaussian mixture models were fitted to the corresponding distribution by employing the Expectation-Maximization (EM) algorithm for mixtures of normal distributions. We first determined the optimal number of Gaussian components (k) using the boot.comp() function from the R mixtools v1.2 package (Benaglia et al. 2009), which performs a parametric bootstrap by producing B bootstrap realizations (replicates) of the likelihood ratio statistic for testing the null hypothesis of a k-component fit versus the alternative hypothesis of a (k+1)-component fit to various mixture models. For this step, we used 1,000 replicates, a significance level of 0.01, and set the maximum number of components to nine. The number of components determined in the previous step was then used to fit a mixture of Gaussian models to the distribution of percent identity values, utilizing the normalmixEM() function from the same package and the following parameters: maxitLJ=LJ1e-30, maxrestartsLJ=LJ1e−3, epsilonLJ=LJ1e−10. The retrieved peaks correspond to percent identity values which equal to the means of the Gaussian mixture components.

## Data Availability Statement

All data generated or analyzed during this study are included in this article and its supplementary information files.

## Abbreviations

NUPT: nuclear plastid DNA sequence
DSB: double-stranded break
NHEJ: non-homologous end joining
SSA: single strand annealing

## Acknowledgements

This work was supported by a “Proyectos I+D Generación de Conocimiento” grant from the Spanish Ministry of Science and Innovation (grant code: PID2020-113277GB-I00) to LCP, and by funds received by the “Sistema de Información Científica de Andalucía” Research Group id BIO359 to LCP. Partially funded by grants PID2019-106758GB-C32 by MCIN/AEI/10.13039/ 501100011033, FEDER “Una manera de hacer Europa” funds, and Junta de Andalucía grant P20-00091 to AS.

## Supplementary Tables

Supplementary Table S1: Distribution of NUPTs across the moringa nuclear genome.

Supplementary Table S2: Distribution of NUPT clusters across the moringa nuclear genome.

